# Maintenance and evolution of individual differences in a prey defence trait examined with a dynamic predator-prey model

**DOI:** 10.1101/2023.12.07.570589

**Authors:** L. Eigentler, K. Reinhold

**Affiliations:** Evolutionary Biology Department, Universität Bielefeld, Konsequenz 45, 33615 Bielefeld, Germany

## Abstract

Predator-prey systems often feature periodic population cycles. In an empirical system with a heritable prey defence trait, ecological oscillations were previously shown to cause evolution of prey defence on the timescale of the population cycles. In this paper, we develop an eco-evolutionary mathematical model comprising partial differential equations to investigate the evolutionary dynamics of prey defence during population cycles. We reveal that ecological population cycles induce evolutionary oscillations of the mean prey defence trait. In contrast to existing continuum modelling frameworks, our model allows for the evolution of individual variability. We show that eco-evolutionary oscillations select for increased individual variability close to the transition from stable to oscillatory states. We also reveal that evolution of prey defence requires both high efficiency and low cost of prey defence and highlight that more information on trade-offs between cost and efficiency is required.

## 1. Introduction

Predator-prey dynamics are a classical example from population dynamics that has been extensively explored through mathematical modelling (see Diz-Pita and Otero-Espinar (2021) for a review). Many theoretical models of predator-prey dynamics exhibit limit cycles. This feature, which is also observed in empirical systems (e.g. Cannon et al. (1976) and Yoshida et al. (2003)), is characterised by time-periodic oscillations of both prey and predator population densities, with the predator density lagging behind the prey density. These cycles are driven by the following mechanism: high prey density leads to an increase of the predator density due to the abundance of the predator’s food source. The resulting high predator density increases the predation pressure on the prey species and leads to a reduction in prey density. This decreased food availability results in a decrease of the predator density and thus a subsequent increase in prey density due to reduced predation pressure. Therefore, both prey and predator species experience a continuous change of their environment on a short timescale. From an evolutionary perspective, this leads to oscillating evolutionary pressures across one single population cycle (Yamamichi 2020).

Population cycles can lead to evolution of prey defence traits on an ecological timescale (Becks et al. 2010; Jones et al. 2009; Yoshida et al. 2003). Prey defence mechanisms reduce the probability of falling victim to predation, for example through improved vigilance (Beauchamp 2015), or camouflage (Skelhorn and Rowe 2016). For example, in an experimental algae-rotifer system (*Chlamydomonas reinhardtii* and *Brachionus calyciflorus*) with a heritable defence trait in the algae population (clumping), increases in predator density led to rapid directional selection for strong prey defence, while during low predator-regimes of the population cycle, directional selection for low prey defence was observed (Becks et al. 2012; Becks et al. 2010). Based on evidence that both prey and predator can adapt to the evolutionary pressures caused by population cycles and vice versa (see Pettorelli et al. (2011), Pettorelli et al. (2015) and Yamamichi (2020) for reviews), a number of theoretical frameworks to investigate these eco-evolutionary dynamics have been developed in recent years (see Velzen et al. (2022) and Yamamichi (2020) for reviews). Eco-evolutionary modelling is not unique to predatory-prey systems but is common in theoretical biology (see Govaert et al. (2018), Klausmeier et al. (2020), Lion (2018) and McPeek (2017) for reviews).

Mathematical modelling of eco-evolutionary dynamics utilise either individual based simulations, or continuum models. Depending on the research questions addressed, continuum modelling of eco-evolutionary predator-prey dynamics typically employs either an adaptive dynamics or a quantitative genetics approach. Quantitative genetics describes the ecological dynamics of an eco-evolutionary system using ordinary or partial differential equations (ODEs/PDEs) whose parameters depend on the mean values of the evolving, normally-distributed traits across the whole population. The evolution of each trait mean is also governed by an ODE, with the rate of change being proportional to the fitness gradient in trait space, and the additive genetic variation in the population. The latter is typically fixed, or a function of the trait mean. The method originates from Lande (1976), Lande (1979), Lande (1981) and Lande (1982), and has more recently been applied to the predator-prey context by Abrams and Matsuda (1997), Cortez (2016), Cortez (2018), Cortez and Patel (2017), McPeek (2017), Schreiber et al. (2011), Velzen and Gaedke (2018) and Velzen et al. (2022).

On the other hand, adaptive dynamics utilises ODEs describing ecological population dynamics to investigate whether rare mutants can invade a resident population with a given strategy under the assumption that ecological dynamics quickly equibrilate. Through the calculation of invasion fitnesses, this method derives conditions for the occurrence of evolutionary stable strategies, or the occurrence of evolutionary branching. Adaptive dynamics has been pioneered by Abrams (2001), Dieckmann and Law (1996), Geritz et al. (1998) and Metz et al. (1995) for simple birth-death processes and has more recently been used in the predator-prey context, for example by Grunert et al. (2021) and MacDonald and Brisson (2022).

While quantitative genetics and adaptive dynamics approaches have many advantages, most and foremost due to their analytical tractability, they do not consider mutation and selection as two separate processes. Moreover, they either assume a separation of timescales between ecological and evolutionary dynamics (adaptive dynamics), or do not allow for the evolution of individual variation itself (quantitative genetics) (Nuismer and Doebeli 2004). The latter is a particularly poignant limitation because individual variation is often used to quantify the occurrence of mutations and is therefore regarded as a crucial necessity for evolution to occur on fast timescales (Barrett and Schluter 2008; Messer and Petrov 2013). For example, under the assumption of fixed individual variation of a prey defence trait, quantitative genetics has shown that predator-prey limit cycles can be caused by large individual variation and that increases in individual variation can increase the phase lag between prey and predator cycles (Cortez 2016). However, this approach could not explore how the ecological dynamics feed back to those of the individual variation and therefore whether these eco-evolutionary dynamics driven by large individual variation are stable over long timescales.

In this paper, we fill this underexplored topic by developing a “trait-diffusion” PDE framework that enables the evolution of individual variation as part of the eco-evolutionary dynamics. Our approach is inspired by recently developed individual based models that have been applied in the contexts of intraspecific competition dynamics among a single population only (Baldauf et al. 2014; Kikuchi and Reinhold 2021; Reinhold et al. 2023), the coevolution of movement strategies in predator-prey systems (Netz et al. 2022), and the evolution of predator perception ranges (Colombo et al. 2019). The idea of “trait diffusion”, in which a trait of interest is considered as an independent variable and individuals can “move” in the “trait space” according to a random walk, has also been applied in continuum models (in which the random walk becomes a diffusion process). In the specific context of a phytoplankton-zooplankton-nutrient system, it was used to show that population limit cycles lead to oscillations in trait mean, and variance and that the trait variance lags behind the trait mean (Merico et al. 2014). Similarly, in the context of an environmental system with resource input pulses it was used to explore evolutionary branching (Tachikawa 2008). The key difference to other eco-evolutionary modelling frameworks, such as adaptive dynamics or quantitative genetics, is that the trait-diffusion approach explicitly splits mutation and selection into two parts; mutation is accounted for by the diffusion of trait values, while selection occurs because the ecological dynamics are considered across the whole trait space.

In this paper we develop a trait-diffusion PDE model for predator-prey dynamics with an evolving prey defence trait that leads to a decrease in predation pressure but is energetically costly. We use the model to explore (i) whether ecological oscillations induce evolutionary oscillations; (ii) under what conditions selection for individual variability occurs over selection for a single strategy of high/low prey defence; and (iii) how eco-evolutionary dynamics are affected by the cost and efficiency of prey defence. Note that we choose prey defence as an example trait. However, we argue that the exploration of other traits, and in particular the coevolution of prey and predator traits are an important topic of future work.

## 2. Methods

We consider an extension of the Rosenzweig-MacArthur predator-prey model that describes predation dynamics alongside the evolution of a prey defence trait (Fig. 1A). The Rosenzweig-MacArthur model is one of the classical ecological predator-prey models featuring population cycles (Rosenzweig and MacArthur 1963). It comprises two ODEs that represent the population dynamics of a prey species *X*(*t*) and a predator species *Y* (*t*) over time *t* ≥ 0, and, in nondimensional form, is given by

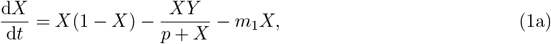

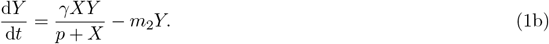

In the model, the prey grows logistically and dies due to predation by the predator, and through natural causes at constant rate *m*_1_ ≥ 0. Often, *m*_1_ = 0 because the logistic growth term implicitly describes prey death. However, we show in Appendix A.2 that it is necessary to choose *m*_1_ ≠ 0 when extending the model to capture evolutionary dynamics. Predation follows a Holling-type II functional response with half-saturation constant *p >* 0. This term represents saturation of the predation pressure caused by a single predator if prey density is high. The predator is assumed to be a specialist predator and therefore its growth is directly proportional (with yield *γ >* 0) to the predation term. Finally, the predator dies naturally at constant rate *m*_2_ *>* 0.

**Figure 1:**
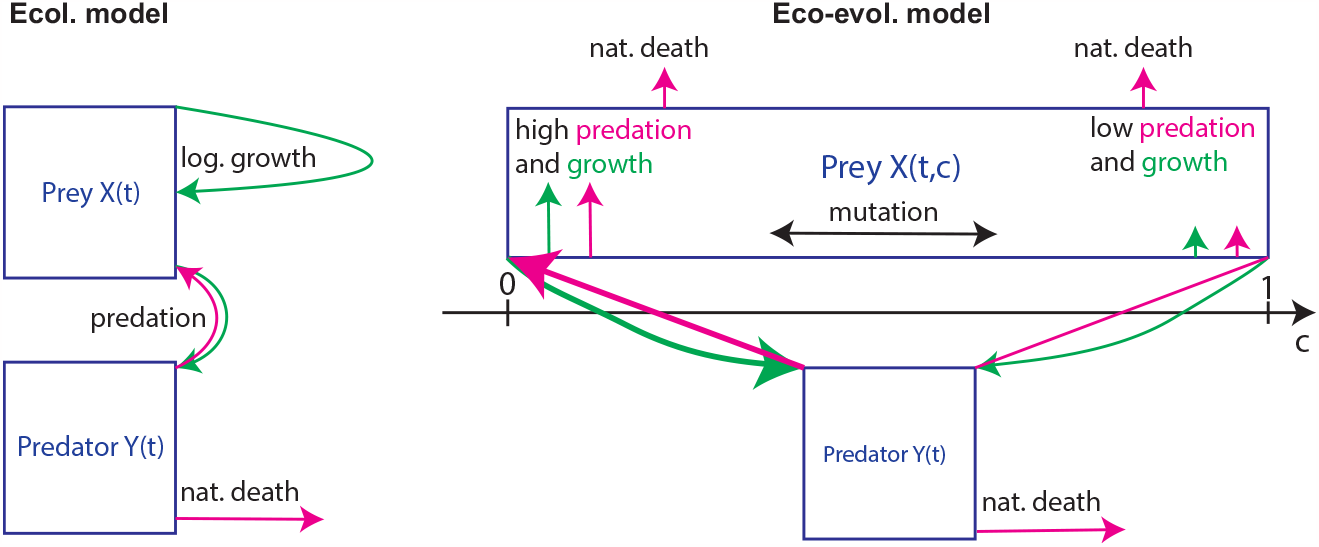
Eco-evolutionary predator-prey dynamics with prey defence. Model schematic for the ecological predator prey model (1) and the eco-evolutionary model (2).

Linear stability analysis (e.g., (Murray 1989)) shows that if natural prey mortality is higher than its maximum growth rate (*m*_1_ *>* 1), then neither prey nor predator can survive. For lower prey mortality (*m*_1_ *<* 1), the model exhibits limit cycles if predator mortality is sufficiently small (*m*_2_ *< γ*(1 − *p* − *m*_1_)*/*(1 + *p* + *m*_1_)), enables constant coexistence of prey and predator for intermediate predator mortality (*γ*(1 − *p* − *m*_1_)*/*(1 + *p* + *m*_1_) *< m*_2_ *< γ*(1 − *m*_1_)*/*(1 − *m*_1_ + *p*)), and causes predator extinction for high predator mortality (*m*_2_ *> γ*(1 − *m*_1_)*/*(1 − *m*_1_ + *p*)).

The Rosenzweig-MacArthur model (1) is a purely ecological model and thus implicitly assumes that there is no individual variation among prey and predator populations and that population traits do not change over time. To investigate how individual variation affects eco-evolutionary dynamics and vice versa, we introduce a prey defence trait as a second independent variable. We denote the prey defence trait by 0 ≤ *c* ≤ 1, where *c* = 0 and *c* = 1 are nondimensional bounds for the minimum and maximum deployment of prey defence, respectively. The lower bound *c* = 0 represents a case of no prey defence, while the upper bound *c* = 1 describes a fully-defended prey under the assumption that prey defence is fully efficient (see function *f*_2_ below). Prey are associated with a “location” in “trait space” and we denote by *X*(*c, t*) the density of prey with trait defence value *c* at time *t*. The predator density is not associated with the prey defence trait and thus remains a function of time only, *Y* (*t*). The prey defence trait is assumed to affect both the predation rate and prey growth rate through a trade off; deploying defensive features uses energy that prey individuals could otherwise use for growth. We define prey growth by

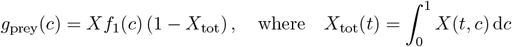

is the total prey density at time *t*, and *f*_1_(*c*) ≥ 0 is a decreasing function. This describes the reduction of growth due to costly prey defence. We assume that increases in the trait *c* also cause a reduction in the predation pressure. For prey with trait *c*, we describe its death due to predation by

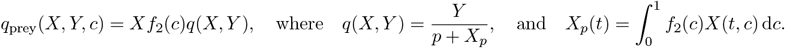

The function *f*_2_(*c*) ≥ 0 is decreasing and thus makes prey with higher trait values less susceptible to predation. As in the ecological model, the maximum per-capita predation rate *q*(*X, Y*) follows a Holling-type II functional response. Note that the denominator is *p* + *X*_*p*_. We refer to Appendix A.1 for more details.

Growth of the predator due to predation is described by integrating the predation term *q*_prey_(*X, Y, c*) over trait space and assuming constant yield *γ >* 0, i.e.,

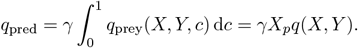

For simplicity, prey are assumed to reproduce asexually and offspring possess the same trait value as their parent. However, prey can change their investment into prey defence through mutation, here modelled by a diffusion term with diffusion coefficient *d >* 0. This yields the eco-evolutionary trait diffusion model

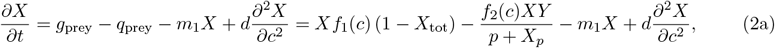

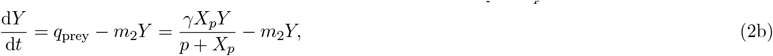

We assume that trait values cannot change beyond the bounds *c* = 0 and *c* = 1 and therefore impose no-flux boundary conditions 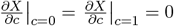. We assume linear responses *f*_1_(*c*) = 1 − *α*_1_*c*, and *f*_2_(*c*) = 1−*α*_2_*c*, where 0 ≤ *α*_1_ ≤ 1 describes the cost of deploying prey defence features and 0 ≤ *α*_2_ ≤ 1 describes the efficiency of the prey defence mechanism.

### 2.1. Model analysis

To numerically solve the model, we discretised the equations in trait space and solved the resulting system of ODEs using Matlab. Simulations were run until an equilibrium state (stable solutions or regular oscillations) was reached. In this paper we refer to *stable solutions* as states in which densities do not change over time to distinguish them from oscillatory states. However, we note that also periodic oscillations are *stable* in a mathematical sense (as perturbations to any stable solution decay over time).

## 3. Results

To address our research questions, we performed a series of numerical simulations of the eco-evolutionary model (2) to construct bifurcation diagrams with respect to three parameters of interest: predator mortality *m*_2_, prey defence cost *α*_1_ and prey defence efficiency *α*_2_. The bifurcation diagrams, visualised in Fig. 2, provide summaries of model outputs that were recorded at equilibrium for given parameters. We report total prey density (i), predator density (ii), mean prey defence trait (iii), and prey defence interquartile range as a measure of individual variability (iv). We report data from constant equilibria in case of stable states (black); for oscillating states, we report the maxima (blue), mean (black), and minima (red) of the oscillating quantities over one period.

**Figure 2:**
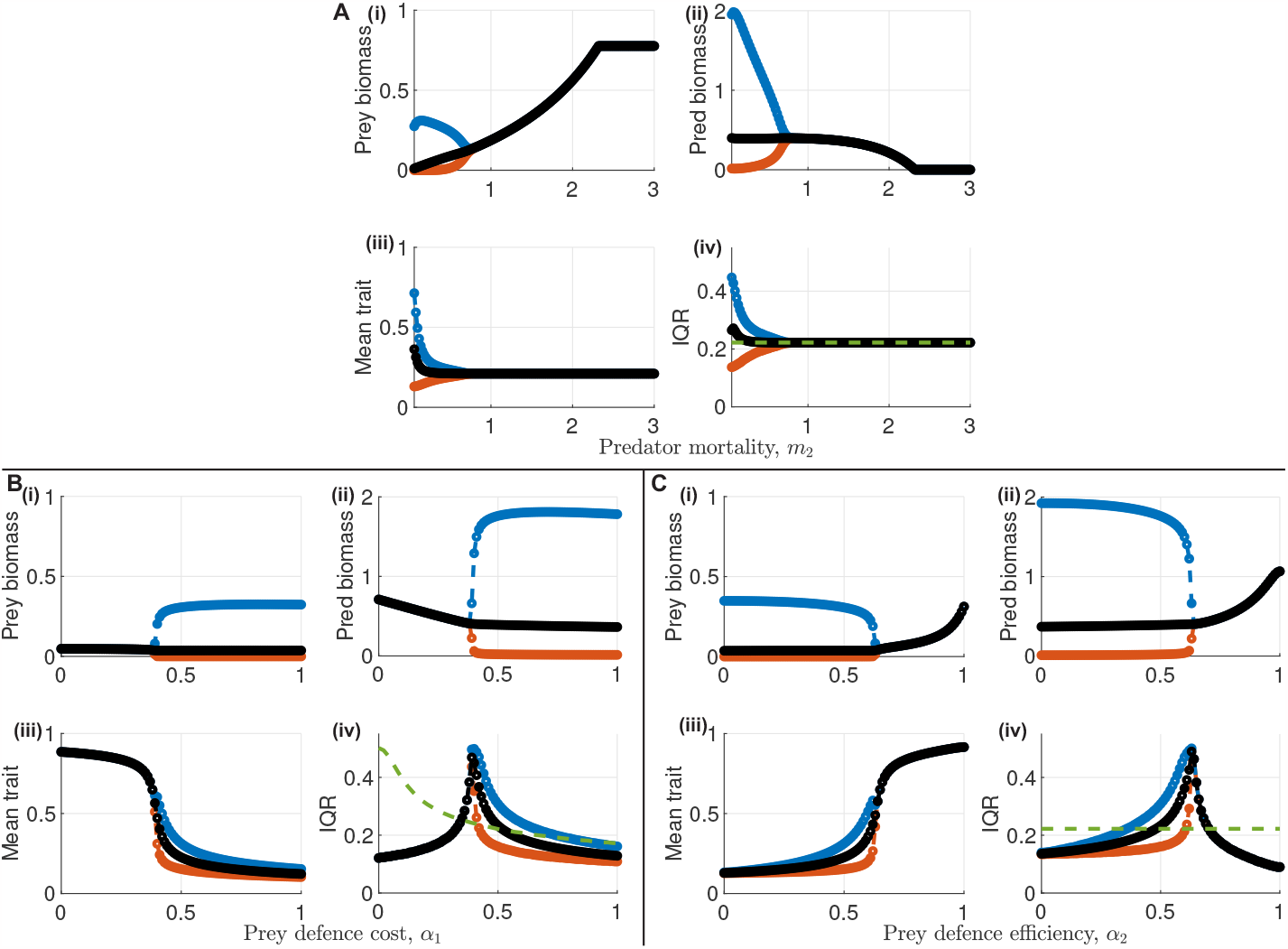
Bifurcation diagrams. **A**: Bifurcation diagram for predator mortality *m*_2_. **B**: Bifurcation diagram for prey defence cost *α*_1_. **C**: Bifurcation diagram for prey defence efficiency *α*_2_. In each panel, parameter values that only show black circles correspond to stable states. Parameter values with black, blue and red circles lead to oscillatory dynamics. Black, blue and red circles show the means, maxima and minima of the quantity of interest across one oscillation, respectively. In (iv) of each panel, the green dashed curve indicates the level of individual variation that occurs due to a mutation-selection balance in the absence of predators. Except the bifurcation parameter, other parameter values are *m*_1_ = 0.2, *m*_2_ = 0.5, *α*_1_ = 0.5, *α*_2_ = 0.5, *p* = 0.5, *γ* = 4, and *d* = 0.001 in each panel.

### 3.1. Ecological oscillations lead to evolutionary oscillations

We first used the bifurcation data to investigate whether the eco-evolutionary model features oscillatory dynamics of predator and prey densities similar to its ecological counterpart (1), and if so, whether these also lead to evolutionary oscillations. Comparing the bifurcation diagram for the predator mortality *m*_2_ (Fig. 2A) with results from the ecological model, we observed that the inclusion of the evolving prey defence trait does not qualitatively affect the bifurcation structure of the system. That is, decreases of predator mortality *m*_2_ cause a transition from a stable predator extinction state to a stable coexistence state to an oscillatory coexistence state (Fig. 2A). Moreover, we found that increases in the prey defence cost *α*_1_, or decreases in prey defence efficiency *α*_2_ cause transitions from stable to oscillatory dynamics (Fig. 2B and Fig. 2C, respectively). We did not observe predator extinction cases in the bifurcation diagrams for prey defence cost and efficiency due to the choice of the predator mortality parameter *m*_2_.

Most importantly, we observed that ecological oscillations of the prey and predator densities also induce oscillations of the mean prey defence trait, and the individual variation within the prey population, independent of which parameter causes a transition from stable to oscillatory states (Fig. 2A-C). An example of an oscillating solution is shown in Fig. 3A-B. We further observed significant changes to the trait distribution across one single period. When predation pressure was low, no investment into prey defence (*c* = 0) was the most common strategy (Fig. 3C). Increases of predation pressure caused the peak of the trait distribution to shift towards higher prey defence trait values (Fig. 3D). Similar to the ecological oscillations, the amplitude of the oscillations of the mean trait and the individual variation increased with decreasing predator mortality *m*_2_ (Fig. 2A). No clear trend on the amplitude of oscillations of trait mean and individual variation was found for changing prey defence cost (Fig. 2B) and efficiency (Fig. 2C). Like in the ecological model, the wavelength of oscillations increased in all cases as parameters were changed to be further from the transition point (Fig. A.3A-C (i)), and the predator cycles lagged slightly behind the prey cycles in all cases (Fig. 3A and Fig. A.3A-C (ii), black). Moreover, there was a further lag between the ecological predator cycles and the evolutionary mean prey defence cycles in all cases (Fig. 3A and Fig. A.3A-C (ii), purple).

**Figure 3:**
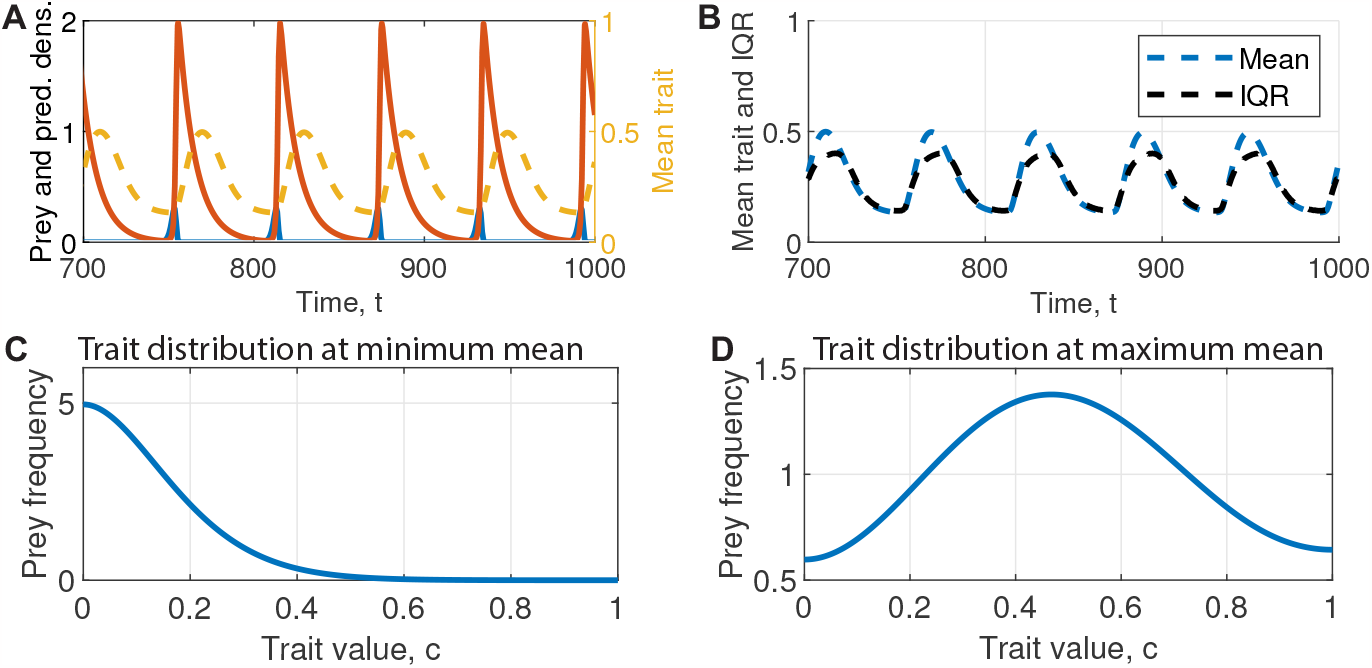
Example oscillatory solution. An oscillatory solution obtained with *m*_1_ = 0.2, *m*_2_ = 0.2, *α*_1_ = *α*_2_ = 0.5, *p* = 0.5, *γ* = 4, *d* = 0.001 is shown. In **A**, the blue curve shows the total prey density, the red curve shows the predator density, and the yellow dashed curve shows the mean trait of the prey population. In **B**, the blue curve shows the mean value of the prey defence trait, and the black curve shows the interquartile range of the prey defence trait distribution. In **C** and **D**, the trait distributions are shown for time points at which the mean trait reaches its minimum (**C**) and maximum (**D**).

### 3.2. Selection for individual variability occurs close to onset of oscillations

We next investigated whether the eco-evolutionary dynamics select for or against individual variability among the prey population. As outlined in Appendix A.2, removing predators from the model (2) leads to selection against prey defence subject to a mutation-selection balance. For all parameter sets shown in the bifurcation diagrams Fig. 2, we quantified how much individual variability occurs in the prey population due to a mutation-selection balance in the absence of predation. For this, we simulated the model with *Y* = 0 and recorded the individual variability at equilibrium. The results are visualised through dashed green curves on the bifurcation diagrams Fig. 2A-C (iii).

Comparisons between the individual variability that occurs in full model simulations with the mutation-selection baseline level revealed that selection for individual variability occurs close to onset of oscillations across all bifurcation diagrams (Fig. 2A-C (iii)). Away from the onset, we observed either lower or similar levels of individual variation when compared with the baseline mutation-selection level.

The detailled response of individual variation differed between parameters. Our bifurcation analysis with respect to predator mortality *m*_2_ revealed that individual variability and mean prey defence trait do not change between the predator extinction and stable coexistence state (Fig. 2A (iii-iv)). This suggests a neutral effect of predator presence on individual prey variability under stable dynamics. However, under oscillatory dynamics, individual variability among prey was on average higher than that caused by the mutation-selection balance, and this average was increasing with decreasing predator mortality (Fig. 2A (iv)). Moreover, the extent of individual variability also oscillated within each eco-evolutionary cycle. It ranged from states in which the distribution was narrower than that caused by the mutation-selection balance (c.f., Fig. A.2B, bottom and Fig. 3C) to states in which individual variation was higher (Fig. 3D). This highlights that while individual variation was selected for on average, selection against individual variation occurred during short timespans within each period of the oscillation. The oscillations of individual variability were in phase with the oscillations in the mean trait value (Fig. 3B).

Changes to the prey defence cost *α*_1_ had a different impact on individual variation (Fig. 2B (iv)). For low costs that resulted in stable dynamics, individual variation was lower than the mutation-selection baseline. Increases of prey defence cost reduced the difference between the baseline and observed variability, eventually leading to a critical threshold at which the variability became larger than the mutation-selection baseline. Increasing prey defence cost increased the variability further until reaching its maximum at approx. IQR= 0.5. Close to this maximum, we also observed the transition from stable to oscillatory states. Beyond the maximum, further increases of prey defence cost caused a decrease of individual variability. Large prey defence costs resulted in individual variability lower than the mutation-selection baseline. Decreases of prey defence cost *α*_2_ had the same qualitatively impact on individual variability as increases to prey defence cost *α*_1_ (Fig. 2C (iv)).

### 3.3. Low prey defence efficiency and high cost prevent prey defence evolution

We next explored how the cost and efficiency of prey defence affected the eco-evolutionary dynamics by further analysing the bifurcation data held in Fig. 2B-C. For stable dynamics, increases in prey defence cost *α*_1_ led to decreases in prey and predator biomass (Fig. 2A (i-ii)). However, changes to prey defence cost did not alter the mean prey and predator biomass in oscillatory dynamics (Fig. 2B (i-ii)). Rather, the amplitude of the oscillations increased in these cases. This suggests that oscillatory dynamics can compensate for increases in prey defence cost.

To investigate this compensatory effect in more detail, we turned to the evolutionary dynamics (Fig. 2B (iii)). The mean prey defence trait was high for low costs of defence and decreased slowly with increasing cost, simultaneously with the biomass decay for stable dynamics. Around the transition to oscillatory dynamics, the mean prey defence trait rapidly decreased from a high level for stable dynamics to a low level for oscillatory dynamics. Further increases in prey defence cost caused further slow decreases in mean prey defence trait, but no further decreases in mean prey and predator biomass, as reported above. Thus, low cost and of prey defence caused selection of high prey defence to counter the predation pressure. However, as higher costs rendered selection for high prey defence unsustainable, it was eco-evolutionary oscillations that enabled the prey to persist in the population and simultaneously facilitated selection for low prey defence. With our assumption that the functional response of prey defence cost to the trait value (*f*_1_(*c*) = 1 − *α*_1_*c*) is identical to that of the efficiency to the trait value (*f*_2_(*c*) = 1 − *α*_2_*c*), we observed qualitatively identical dynamics for increasing prey defence cost *α*_1_ and decreasing prey defence efficiency *α*_2_ (c.f., Fig. 2B and C).

We next tested how the switch between stable and oscillatory dynamics and the associated switch between evolution towards high and low prey defence is affected by simultaneous changes to prey defence cost *α*_1_ and efficiency *α*_2_. For this, we performed numerical simulations on a grid of *α*_1_ and *α*_2_, and determined the location of the switch in this parameter plane (Fig. 4; black).

**Figure 4:**
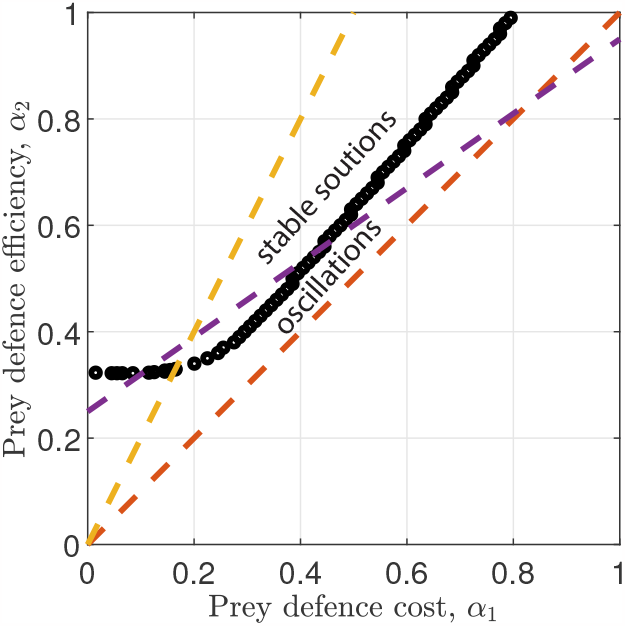
Switch between stable and oscillatory dynamics under simultaneous variation of prey defence cost and efficiency. The location of the switch between oscillatory and stable dynamics is shown in the parameter plane spanned by prey defence cost *α*_1_ and the prey defence efficiency *α*_2_. Note that only this switch is shown in the plot; no information on switches to extinction states are displayed. The dashed lines show different possible linear trade-offs between *α*_1_ and *α*_2_. The parameter values used are *m*_1_ = 0.2, *m*_2_ = 0.2, *p* = 0.5, *γ* = 4, *d* = 0.001.

This scan of the parameter plane revealed that the switch from oscillatory dynamics to stable dynamics caused by decreases in prey defence cost *α*_1_ (reported above) could only occur if prey defence efficiency *α*_2_ was sufficiently high. Vice versa, switches from oscillatory to stable dynamics caused by increases in prey defence efficiency *α*_2_ could only occur if prey defence cost *α*_1_ was sufficiently low. Thus, high prey defence cost or low prey defence efficiency always led to oscillatory states (Fig. 4). As reported above, oscillatory states were always characterised by evolution of low prey defence. Thus, low prey defence efficiency *α*_2_ prevented the evolution of prey defence even in cases when prey defence cost was low. Similarly, high prey defence costs *α*_1_ prevented the evolution of prey defence even when its efficiency was high.

This observation highlights that switches between stable and oscillatory states depend on the trade-off between prey defence cost *α*_1_ and efficiency *α*_2_. Fig. 4 shows three possible linear trade-offs *α*_1_ = *a* + *bα*_2_ (dashed lines). Even such a simple form of the trade-off can, depending on the slope and intersect of the linear relation, lead to either no (red in Fig. 4), one (yellow), or two (purple) switches along the trade-off curves, with other, non-linear trade-offs possibly leading to even more complex dynamics.

## 4. Discussion

In this paper we used a trait-diffusion model to reveal the eco-evolutionary feedbacks in predator-prey systems with an evolving prey defence trait. Most importantly, we revealed that (i) ecological oscillations induce evolutionary oscillations of both mean trait and individual variability; (ii) individual variability in prey defence is selected for close to the onset of oscillations; and (iii) evolution of prey defence only occurs if cost is low and efficiency is high. Significantly, we highlight that exploring the impacts of changes to the trade-off between prey defence cost and efficiency and linking these to empirical data is an important challenge for future work, and that any future modelling approaches should use frameworks that allow for the evolution of individual variability.

### Ecological oscillations induce evolutionary cycles

The occurrence of periodic oscillations of prey and predator densities is a well-known phenomenon in both empirical and theoretical predator-prey systems (Blasius et al. 2019; Diz-Pita and Otero-Espinar 2021). Our results highlight that these ecological oscillations also induce periodic dynamics of a prey defence trait that evolves on the same timescale as the ecological dynamics take place (e.g., Fig. 3). Moreover, oscillations of both mean trait and individual variation lag behind the predator (up to a quarter period) and prey (up to half a period) densities (Fig. A.3); this is in addition to the lag between prey and predator oscillations which is well documented in both empirical and theoretical literature (Blasius et al. 2019; Diz-Pita and Otero-Espinar 2021; Velzen and Gaedke 2018). Intuitively, the evolutionary oscillations occur as follows. Starting from a state of high prey density and low predator density in the ecological cycle, increases in predator density increase predation pressures. Therefore selection for higher investment in prey defence occurs. However, high investment into prey defence causes a reduction in average predator net growth. As the predator density decreases beyond its peak, high investment into prey defence becomes energetically unsustainable. Thus, selection for low prey defence occurs. This increases predator net growth and thus closes the loop. Phase shifts between ecological and evolutionary oscillations have previously been reported from an empirical system involving *C. reinhardtii* (prey) with the ability to clump (prey defence) and *B. caluciflorus* (predator) (Becks et al. 2012). We find that the high adaptability of prey to predation pressure allows the prey population to persist even if the cost of prey defence *α*_1_ is high or its efficiency *α*_2_ is low (Fig. 2B and C, respectively). We note that the observation that ecological oscillations induce cycles in mean trait values (prey defence or others) has previously been reported from eco-evolutionary predator-prey models in which individual variability was a fixed input parameter (Abrams and Matsuda 1997; Cortez 2016; Cortez 2018; Cortez and Patel 2017; Schreiber et al. 2011). By contrast, our results reveal that individual variation also oscillates provided individual variation is considered as an emergent feature rather than a fixed input. This result is in agreement with the work by Merico et al. 2014 who used a trait diffusion model to describe the specific case of zooplankton-phytoplankton interactions with an edibility trait in the phytoplankton population.

### Selection for individual variability occurs close to onset of oscillations

Our results further highlight that individual variability in prey defence is selected for under a wide range of parameter values close to the onset of eco-evolutionary cycles (Fig. 2A-C (iv)). While some individual variability occurs solely through a mutation-selection balance, we observed larger individual variability for intermediate prey defence costs, intermediate prey defence efficiencies and low predator mortalities (Fig. 2). The common factor among the results for these parameter space scans is that selection for individual variability always occurred close to the transition between stable and oscillatory dynamics. In this context, it is important to note that oscillatory dynamics far from the transition point are likely of no biological relevance. Far from onset, oscillations in our deterministic model feature extremely low values of both biomass densities during troughs of the oscillations. In stochastic empirical systems, however, it is inevitable that periodically occurring times of extremely low densities eventually lead to an extinction event caused by a environmental stochasticity (Alkhayuon et al. 2023). On the other hand, stable dynamics far from the transition to oscillations possess biological relevance. Thus, we conclude that biologically relevant oscillatory dynamics lead to selection for individual variability, while stable dynamics can feature a range of evolutionary outcomes depending on parameter values.

Often, individual variability is attributed to negative frequency dependent selection (Baldauf et al. 2014; Kikuchi and Reinhold 2021; Wolf et al. 2007) or exogenous forces that cause temporal and/or spatial fluctuations in selection (Reinhold 2000; Svardal et al. 2015). We have not included any of these features in our model systems, and thus conclude that individual variability is caused by other processes. Rather, we present evidence that individual variability occurs for parameters that lead to a balance between costs and benefits of prey defence, independent of the trait value. Such a balance is a known consequence of dynamics in which growth rates trade-off with mortality rates (here through protection from predation) (Mangel and Stamps 2001). The closer the system is to such a balance, the less difference in individual net growth rate (fitness) exists between different genotypes along the prey defence trait axis. In our model, individual net growth rate for one genotype is *g*(*X, Y* ; *c*)*/X*, where *g*(*X, Y* ; *c*) is the right hand side of (2a). In Fig. 5A, we visualised the individual net growth rates of genotypes for three different parameter values that resulted in low, medium, and high individual variability (the trait distributions are shown in Fig. 5B) when constructing the bifurcation diagram shown in Fig. 2B. These data highlight that higher individual variability occurred when there was less growth rate differences between individual genotypes. This can be interpreted in terms of a mutation-selection balance. The lower the fitness differences along the trade space, the less importance is given to selection. Since mutation is not directional, this leads to increased individual variation.

**Figure 5:**
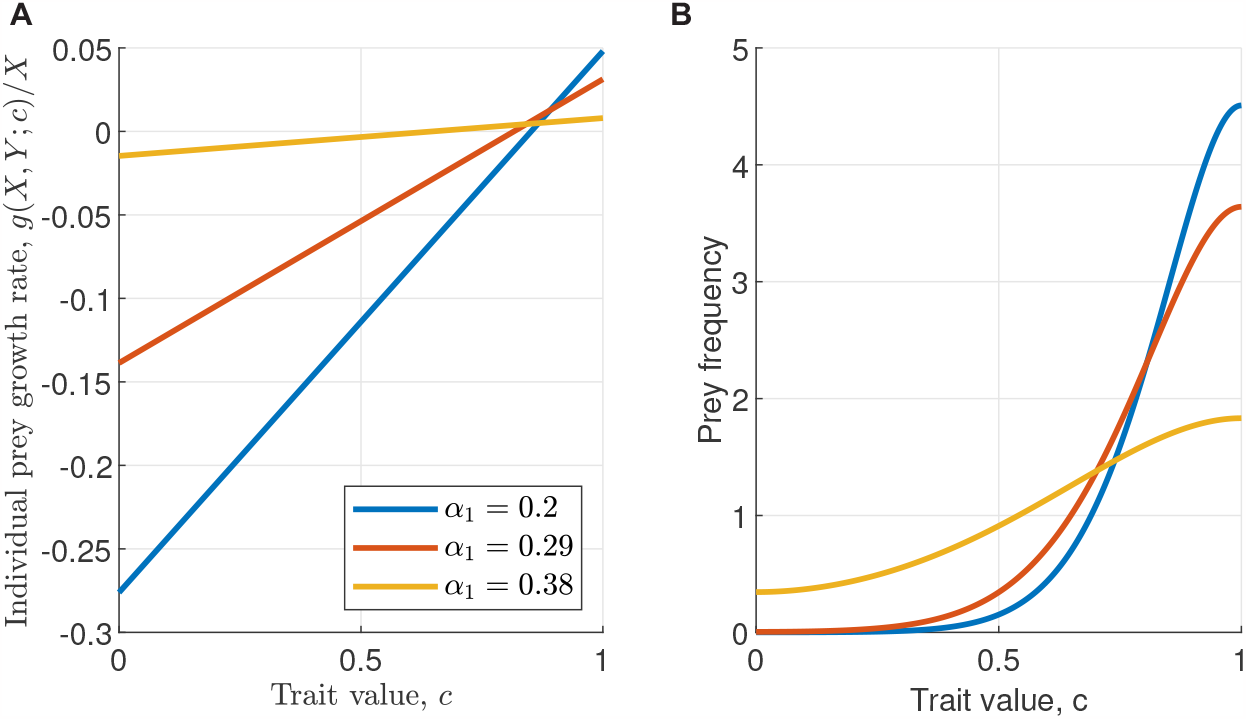
Individual growth rate along the prey defence trait axis. **A**: The individual prey growth rates *g*(*X, Y* ; *c*)*/X* for all prey defence genotypes are shown for three different parameter values. All parameter values correspond to choices in Fig. 2B and represent cases that resulted in high (yellow), medium (red), and low (blue) individual variability among prey. **B**: The trait distributions for the three cases in part A are shown.

### Future work should focus on the trade-off between prey defence cost and efficiency

We remark the importance of exploring in more detail the impact of trade-offs between prey defence cost and efficiency in future work. Our results highlight that, under the assumption of linear relationships (through the functions *f*_1_(*c*) and *f*_2_(*c*)) between trait values *c* and their impact on prey growth and susceptibility to predation, stable solutions can only occur if prey defence is both efficient (high *α*_2_) and cheap (low *α*_1_) (Fig. 4). In other words, we found that low costs could not compensate for low efficiencies and high efficiencies could not compensate for high costs. Exploring the precise impact of different trade-offs between prey defence cost and efficiency, and different functional responses of prey defence cost and efficiency to trait values was not the goal of this paper. However, results from this paper emphasise that such an exploration is a important topic of future work in order to predict switches between stable and oscillatory dynamics and associated evolutionary consequences. Indeed, there is plenty of evidence in empirical data that trade-offs between prey defence cost and efficiency exist. For example, cluster (palmella) formation by algae of *C. reinhardtii* that reduces predation by the rotifer species *B. calyciflorus* was shown to reduce the growth rate of *C. reinhardtii* in the absence of predation (Becks et al. 2010), with similar results existing for predator-prey dynamics between *Chlorella vulgaris* and *B. calyciflorus* (Meyer et al. 2006). Further, plants which develop defensive traits that reduce herbivory have been shown to be more susceptible to plant pathogens (Felton et al. 1999; Strauss et al. 2002; Thaler et al. 1999), or reductions in pollination (Strauss et al. 2002; Strauss et al. 1999). Moreover, several previous empirical studies highlighted that prey defence cost can change over time due to dependence on environmental factors (Kraaijeveld and Godfray 1997; Yoshida et al. 2004). Combined, this emphasises the need to develop a comprehensive understanding of the impact of prey defence cost-efficiency trade-offs.

### Comments on model choice

Finally, we highlight the importance of using a modelling approach that allows for the evolution of individual variability. Previous theoretical studies that have opted for a continuum modelling approach (Merico et al. 2014 being a notable exception) of eco-evolutionary predator-prey dynamics (considering prey defence and other traits) have typically employed quantitative genetics frameworks (e.g., Abrams and Matsuda 1997; Cortez 2016; Cortez 2018; Cortez and Patel 2017; McPeek 2017; Schreiber et al. 2011; Velzen and Gaedke 2018; Velzen et al. 2022). While this approach often results in model systems that are analytically tractable, it requires the assumption that trait distributions are fixed to be normal with either fixed variances that cannot evolve or higher order moment closures. The results presented in this paper highlight that these assumptions present major limitations. We therefore argue that future theoretical approaches of eco-evolutionary dynamics (not necessarily limited to predator-prey systems) should focus more on model types that allow for the evolution of both the shapes of trait distributions and their variances. In this paper, we employed a “trait-diffusion” approach that describes mutation events as a random walk process and allows selection to occur explicitly through the ecological dynamics. Similar freedom (in terms of the evolution of trait distributions) is afforded by individual based models (e.g. Baldauf et al. 2014; Colombo et al. 2019; Kikuchi and Reinhold 2021; Netz et al. 2022; Reinhold et al. 2023), and an extension of the quantitative genetics framework, termed “oligomorphic dynamics”, that constructs trait distributions as the sum of an arbitrarily large number of normal distributions (Lion et al. 2022; Lion et al. 2023; Sasaki and Dieckmann 2011).

## Acknowledgments

The authors thank Xander O’Neill (Heriot-Watt University, UK), Meike Wittmann (Universiät Bielefeld, Germany) and David W. Kikuchi (Oregon State University, US) for helpful comments and discussions on an earlier draft of the paper.

## Data availability

Computational code used to obtain the results presented in this paper has been deposited in a Github repository which has been archived through Zenodo (Eigentler and Reinhold 2023).

## Author contributions

The author statement uses CRediT (Contributor Roles Taxonomy). Lukas Eigentler: Conceptualization, Methodology, Software, Formal analysis, Investigation, Data Curation, Writing - Original Draft, Writing - Review & Editing, Visualization, Project administration, Funding acquisition. Klaus Reinhold: Conceptualization, Writing - Review & Editing, Project administration, Funding acquisition.

### A. Appendix

#### A.1. Holling-type II functional response in the eco-evolutionary model

The maximum per-capita predation rate *q*(*X, Y*) in the eco-evolutionary model follows a Holling-type II functional response. The denominator of the term is *p* + *X*_*p*_, rather than *p* + *X*_tot_. That is, saturation of the predation pressure exerted by a single predator depends on the abundance of prey and its trait distribution, rather than just the prey abundance. This means that for fixed total prey density and given prey defence trait value, predation pressure on prey individuals depends on the proportion of well-defended prey in the population. More well-defended prey in the population lead to higher predation pressure for each prey individual compared with a state in which there are less well-defended prey; this is visualised in Fig. A.1B in which the per capita predation pressure for each trait value is compared for two different trait distributions. The dependence on the prey trait distribution represents either the predators’ knowledge of the prey’s defence trait, or the predators’ persistence in continuing predation on other individuals after failed attempts on well-defended prey.

#### A.2. Explicit modelling of prey mortality is essential to capture realistic evolutionary dynamics

Before using the model to address questions on the evolution of prey defence, we discuss the inclusion of the prey mortality term −*m*_1_*X* in (2). Justification is required because ecological modelling of predator-prey dynamics using Rosenzweig-MacArthur-type models typically do not explicitly model prey death for reasons other than predation (i.e., set *m*_1_ = 0). The rationale is that the logistic growth term *X*(1 − *X*) in (1) describes the *net growth* of the prey population in the absence of predation and thus includes any predator-independent causes of death. For the purpose of modelling ecological dynamics, this implicit approach of describing birth and death processes is a reasonable assumption because all prey individuals are assumed to be identical.

By contrast, it is essential to explicitly model predator-independent (and therefore trait-independent) prey mortality when describing eco-evolutionary dynamics, as the following simulations of the eco-evolutionary model show. Consider a setting without any predators. In this case, the eco-evolutionary model reduces to just the prey dynamics in the absence of predation ((2a) with *Y* = 0). Intuitively, any theoretical framework successfully describing the evolution of prey defence that trades-off with prey growth should predict directional selection towards no prey defence in the absence of predators. However, in the absence of predators (*Y* = 0) and predator-independent prey mortality (*m*_1_ = 0), the eco-evolutionary model (2) predicts evolution towards an equilibrium with a uniform distribution of the prey defence trait, i.e. a state in which all possible prey defence strategies occur at the same frequency (Fig. A.2A). Directional selection of no prey defence only occurs provided predator-independent prey mortality is considered (*m*_1_≠ 0). In this case, the prey population evolves to a state with small mean prey defence trait value (Fig. A.2B). The remaining deviation from zero occurs due to a mutation-selection balance. Unsurprisingly, the size of the deviation depends on the mutation rate *d* (Fig. A.4). We use *d* = 0.001 throughout the paper but note that any statements on individual variation depend quantitatively on the mutation rate. Note also that the inclusion of predator-independent prey mortality reduces the prey density at equilibrium.

**Figure A.1:**
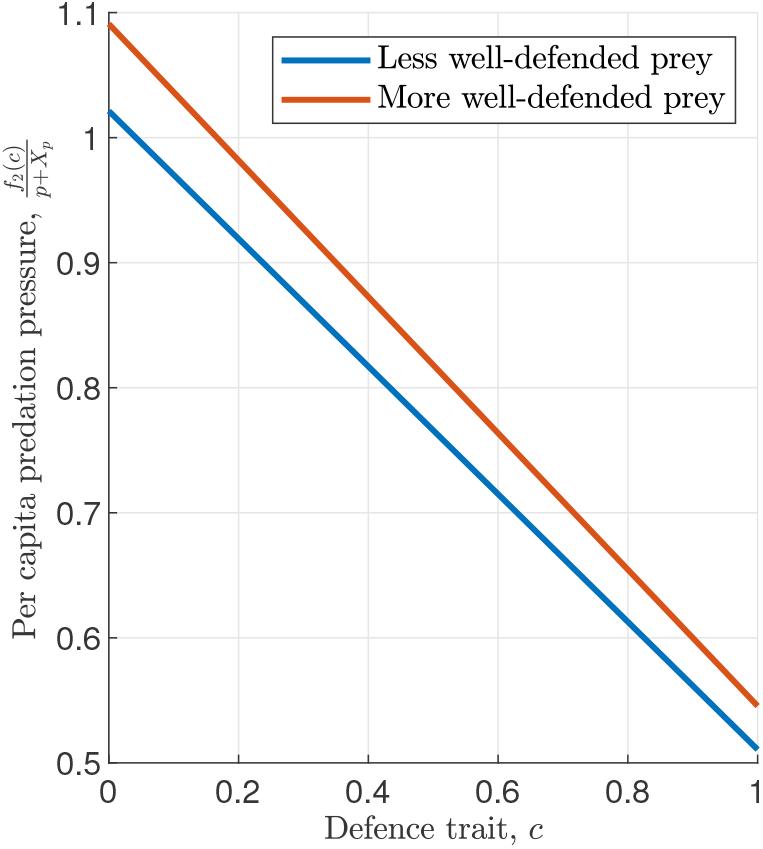
Higher relative abundance of well-defended prey increase predation pressure on all individuals. Per capita predation pressure under fixed total prey density and different prey trait distributions. The red curve represents a case in which there are more well-defended prey than in the case represented by the blue curve. The probability density functions are *F* (*X*) = 0.75 + 0.5*c* and *F* (*X*) = 2*c*, for blue and red, respectively. Higher abundances of well-defended prey increase the predation pressure for each single individual due to predators’ knowledge of prey defence traits or their persistence to continue predation after a failed attempt on well-defended prey.

**Figure A.2:**
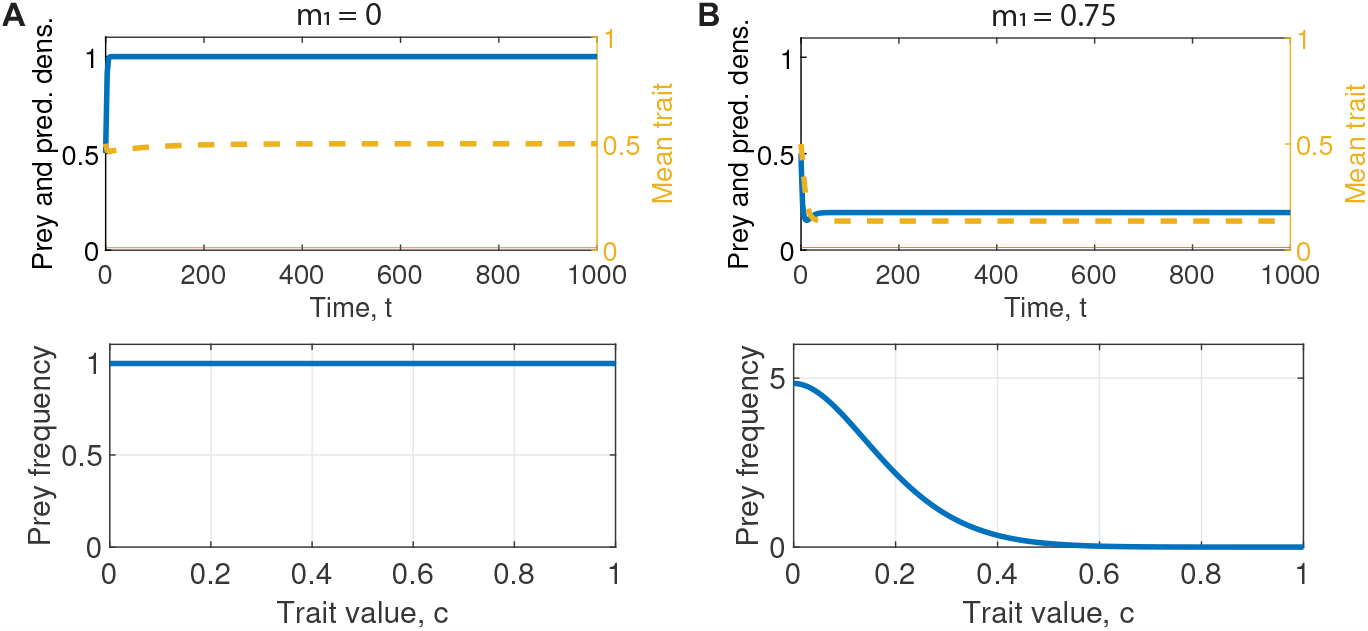
Explicit description of predator-independent prey mortality is required to capture realistic evolution of prey defence. A comparison between the exclusion (A) and inclusion (B) of predator-independent prey mortality is shown for the eco-evolutionary dynamics of a prey population in the absence of the predator. In both A and B, the top panel shows the total prey (blue) density, its mean trait value (yellow), and the predator density (red; here kept at zero). The bottom panel shows the trait distribution at the end of the simulation. The predator-independent prey mortality is *m*_1_ = 0 in A, and *m*_1_ = 0.75 in B. Other parameter values are *d* = 0.001, *α*_1_ = 0.5.

This highlights that explicit description of predator- and trait-independent prey mortality is necessary to capture realistic eco-evolutionary dynamics. We therefore use *m*_1_≠ 0 in the remainder of the paper.

#### A.3. Other supplemental figures

**Figure A.3:**
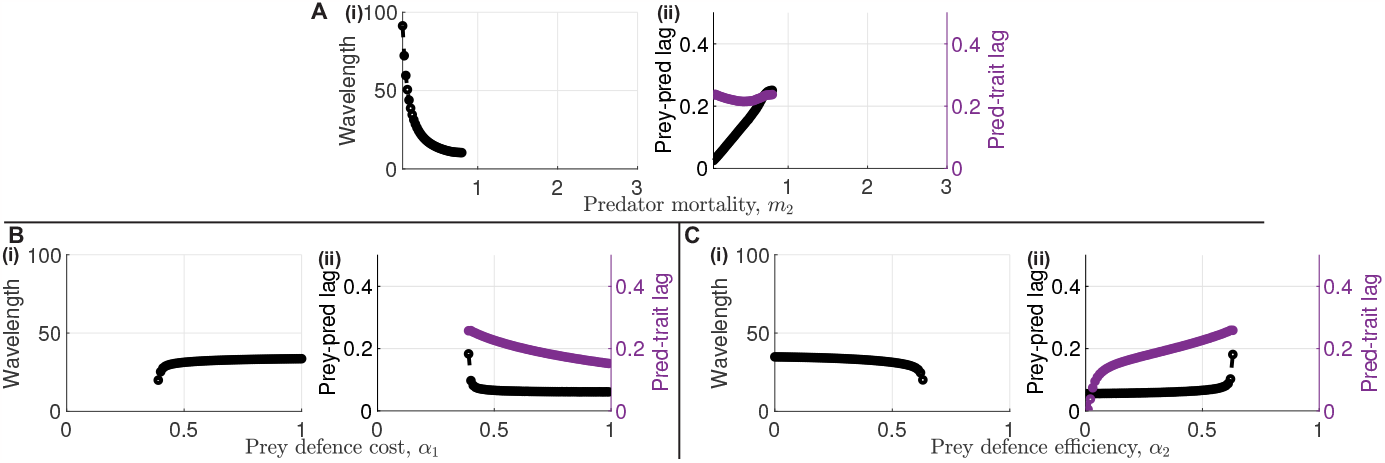
Wavelengths and phase-lags. **A**: Wavelength and phase-lag for changes to predator mortality *m*_2_. **B**: Wavelength and phase-lag for changes to prey defence cost *α*_1_. **C**: Wavelength and phase-lag for changes to prey defence efficiency *α*_2_. In (i) of each panel, the wavelength is shown for parameters that lead to oscillations in the bifurcation diagram Fig. 2. Similarly, in (ii) of each panel, the lag between prey and predator oscillations (black) and lag between predator and prey defence trait oscillations (purple) are shown for parameters that lead to oscillations in the bifurcation diagram Fig. 2. Except the bifurcation parameter given on the *x* axis, other parameter values are *m*_1_ = 0.2, *m*_2_ = 0.5, *α*_1_ = 0.5, *α*_2_ = 0.5, *p* = 0.5, *γ* = 4, and *d* = 0.001 in each panel.

**Figure A.4:**
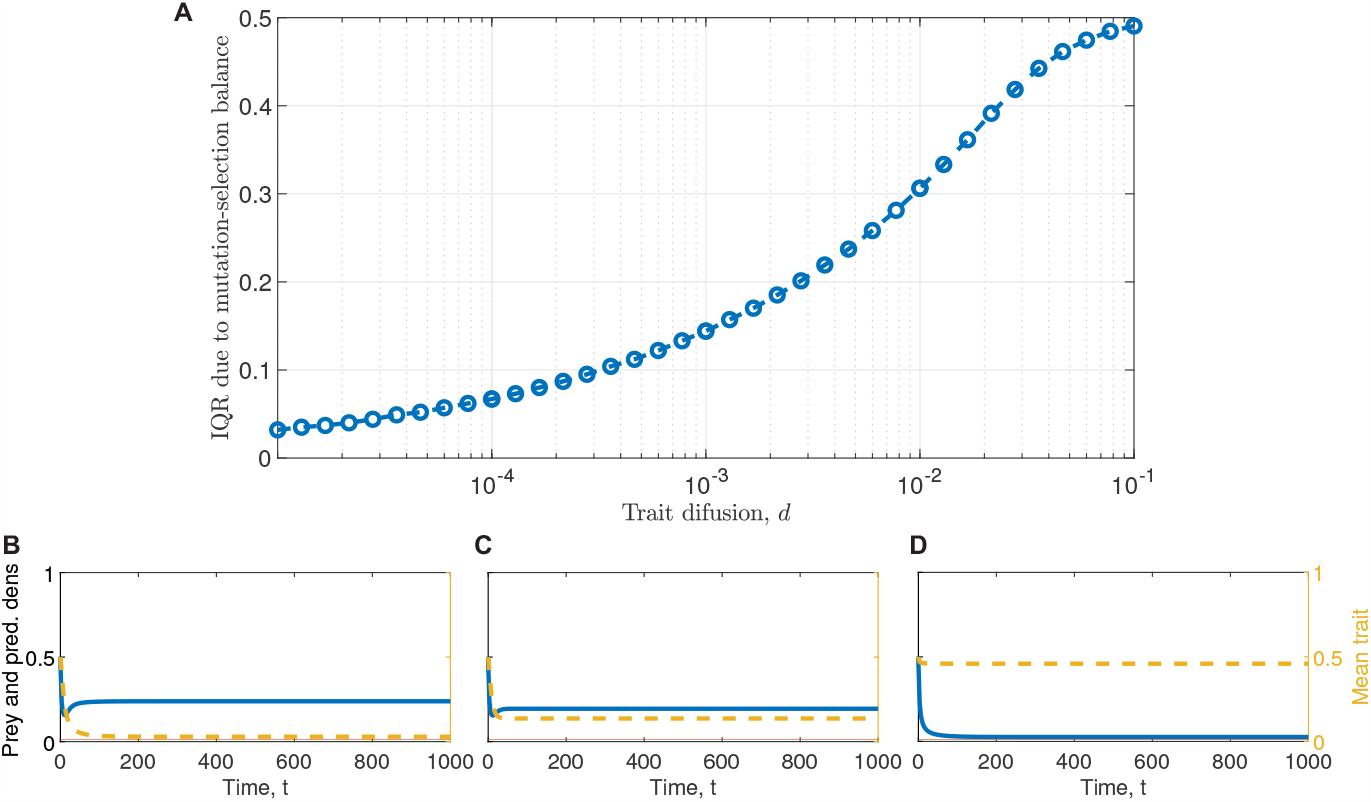
Trait diffusion strength and mutation-selection balance. **A**: This figure visualises the interquartile range of the prey defence trait that occurs in the absence of predators (*Y* = 0) for a range of different trait diffusion strengths *d*. Thus, it represents the amount of individual variability that is due to a mutation-selection balance. **B-D**: Example solutions for *d* = 10^−5^ (B), *d* = 10^−3^ (C), and *d* = 10^−1^ (D). The blue curve shows the total prey density, the red curve shows the predator density (here set to zero), and the yellow dashed curve shows the mean trait of the prey population. Other parameter values are *m*_1_ = 0.75 and *α*_1_ = 0.5.

